# Universal Design Principle to Enhance Enzymatic Activity using the Substrate Affinity

**DOI:** 10.1101/2023.02.01.526728

**Authors:** Hideshi Ooka, Yoko Chiba, Ryuhei Nakamura

## Abstract

Design principles to improve enzymatic activity are essential to promote energy-material conversion using biological systems. For more than a century, the Michaelis-Menten equation has provided a fundamental framework of enzymatic activity. However, there is still no concrete guideline on how the parameters should be optimized to enhance enzymatic activity. Here, we demonstrate that tuning the Michaelis-Menten constant (*K*_*m*_) to the substrate concentration (*S*) maximizes enzymatic activity. This guideline (*K*_*m*_ = *S*) was obtained by applying the Brønsted (Bell)-Evans-Polanyi (BEP) principle of heterogeneous catalysis to the Michaelis-Menten equation, and is robust even with mechanistic deviations such as reverse reactions and inhibition. Furthermore, *K*_*m*_ and *S* are consistent to within an order of magnitude over an experimental dataset of approximately 1000 wild-type enzymes, suggesting that even natural selection follows this principle. The concept of an optimum *K*_*m*_ offers the first quantitative guideline towards improving enzymatic activity which can be used for highthroughput enzyme screening.

## MAIN TEXT

### Introduction

Enzymes are responsible for catalysis in virtually all biological systems,^[1,2]^ and a rational framework to improve their activity is critical to promote biotechnological applications. Since the early 20^th^ century, a reaction mechanism where the enzyme first binds to the substrate (E+S → ES) before releasing the product (ES → P) has been used as the conceptual basis to understand enzyme catalysis (Scheme 1).^[3-6]^ The reaction rate of this mechanism is given by the Michaelis-Menten equation:

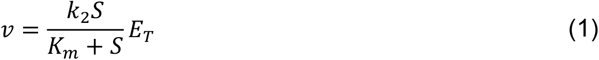

Here, the reaction rate (*v*) is expressed as a function of a rate constant (*k*_2_), the Michaelis-Menten constant (*K*_*m*_), and the substrate (*S*) and enzyme (*E*_*T*_) concentrations. *K*_*m*_ can be interpreted as a quasi-equilibrium constant for the formation of the enzyme-substrate complex, defined as:

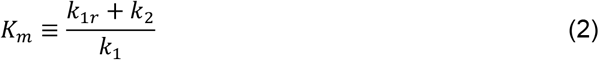

with rate constants defined based on the mechanism shown in Scheme 1. *k*_2_ is the rate constant for releasing the product from the enzyme-substrate complex (ES → P), routinely expressed as *k*_*cat*_ in the enzymology literature. These parameters are experimentally accessible by fitting the theoretical rate law (Eq. (1)) with experimental data^[7-10]^ and are subsequently registered in databases such as BRENDA^[11]^ and Sabio-RK.^[12]^ In principle, the accumulated data may help rationalize and improve the activity of existing enzymes.

**Scheme 1.**
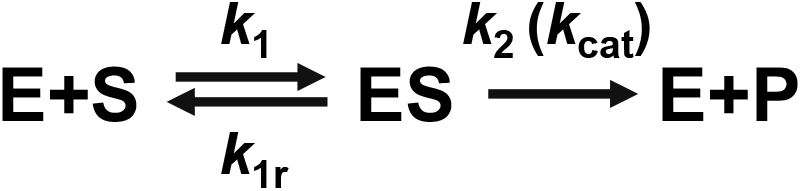
The standard reaction mechanism of enzyme catalysis.

However, there is no concrete understanding on how these parameters influence enzymatic activity. For example, increasing *k*_2_ may enhance activity according to Eq. (1), or diminish it due to a larger *K*_*m*_ (Eq. (2)).^[13]^ Thus, the mutual dependence between *k*2 and *K*_*m*_ complicates their influence on the enzymatic activity ( *v* ), hindering the rational design of enzymes towards biotechnological applications such as the synthesis of commodity chemicals,^[14]^ antibiotics,^[15]^ or pharmaceuticals,^[16]^ increasing the nutritional content of crops,^[17]^ and restoring the environment.^[18]^

In this study, we analyzed the Michaelis-Menten equation to clarify the relationship between the enzyme-substrate affinity (*K*_*m*_) and the activity (*v*). The key ingredient of our mathematical analysis is the Brønsted (Bell)-Evans-Polanyi (BEP) relationship,^[19-23]^ which models the activation barrier as a function of the driving force. This is a well-known concept in heterogeneous catalysis, and in conjunction with the Arrhenius equation,^[24]^ can be used to evaluate the mutual dependence between *k*_2_ and *K*_*m*_ to quantitatively. This allowed us to calculate the optimum value of *K*_*m*_ required to maximize enzymatic activity (*v*), a finding which is supported by our bioinformatic analysis of approximately 1000 wild-type enzymes.

## Results

### Construction of the Thermodynamic Model

In principle, an ideal enzyme with low *K*_*m*_ and large *k*2 can be realized if both *k*1 and *k*2 are increased simultaneously. However, this is physically unrealistic, because the driving force which can be allocated to *k*_1_ and *k*_2_ is limited by the free energy change of the entire reaction. Within this thermodynamic context, maximum activity is realized by optimizing the distribution of the total driving force between the first (E +S → ES) and second (ES → P) steps shown in Scheme 1. To quantitatively evaluate the relationship between the driving force and the activity, we have used the BEP relationship^[19-23]^ to convert driving forces ( Δ*G* ) into activation barriers ( *Ea* ), and the Arrhenius^[24]^ equation to convert activation barriers to rate constants.

The thermodynamic model which served as the basis of our calculations is shown in Fig. 1. In a classical Michaelis-Menten reaction, the enzyme and substrate first form an enzyme-substrate complex (E+S → ES) before producing the product in the second step (ES → P). This mechanism is conceptually similar to reactions that occur on a heterogeneous catalyst surface, where the substrate molecule first binds to the catalyst surface before being converted into the product.^[19-23]^ The Gibbs free energies for the formation of the enzyme-substrate complex and the product are denoted as Δ*G*_1_ and Δ*G*_2_, respectively. By definition, their sum must equal the total free energy change of the reaction Δ*GT*:

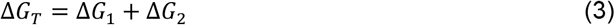

**Figure 1.**
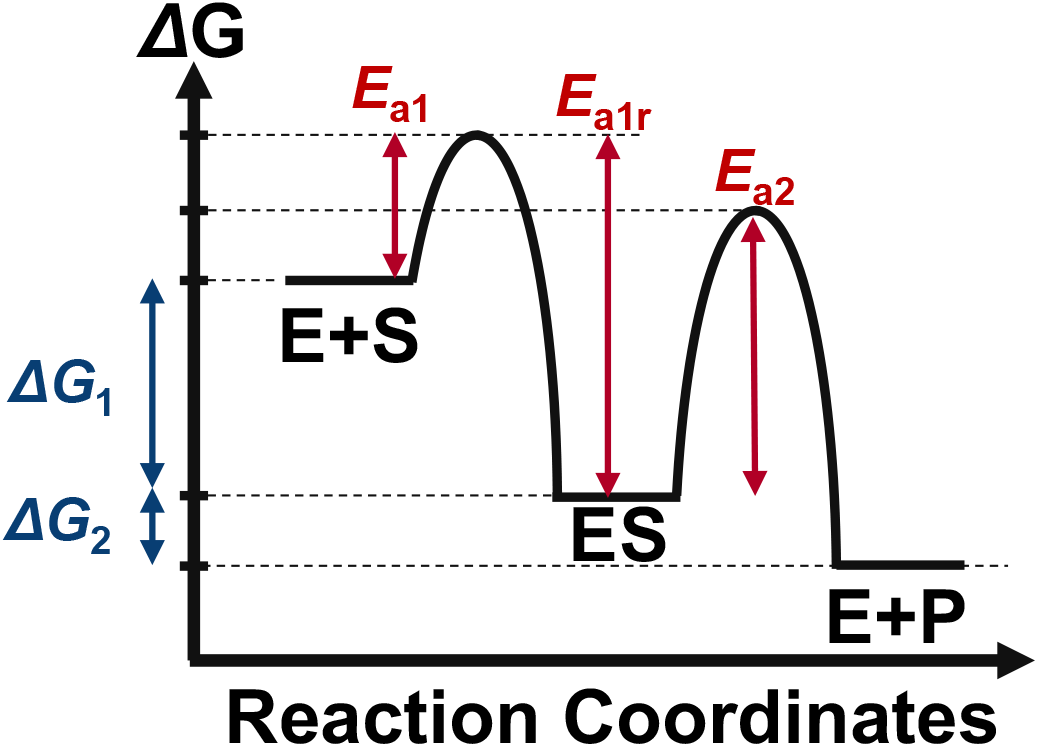
The free energy landscape corresponding to the mechanism shown in Scheme 1. Each reaction in the mechanism is labeled by its corresponding rate constant. The free energy landscape below indicates the free energy changes (Δ*G*_1_, Δ*G*_2_) and activation barriers (*E*_*a*1_, *E*_*a*1*r*_, *E*_*a*2_).

## Reaction Coordinates

From these thermodynamic constraints, we will use the BEP relationship^[19-23]^ to obtain activation barriers (*Ea*), and then the Arrhenius^[24]^ equation to obtain rate constants, which ultimately yields quantitative insight on the relationship between *k*_1_, *k*_2_, and *K*_*m*_. Based on the BEP relationship, the activation barrier corresponding to *k*_1_ can be written as:

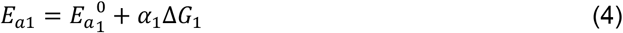

where 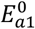represent the activation barriers when the elementary reaction is in equilibrium (Δ*G*_1_ = 0). They are positive constants which reflect the inherent favorability of this elementary step. α_1_ is a positive constant coefficient which indicates the sensitivity of the activation barrier with respect to the driving force. Recently, Kari et al have shown that fungal cellulases indeed satisfy such linear free energy relationships between the activation barrier and the driving force.^[25]^ Next, activation barriers can be converted to rate constants based on the Arrhenius equation^[24]^ as follows:

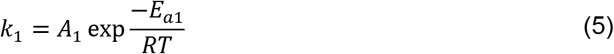

Here, *A*_1_ is a pre-exponential factor, and *R* and *T* are the gas constant and absolute temperature, respectively. Using Eqs. (4) and (5), *k*_1_ can be expressed as:

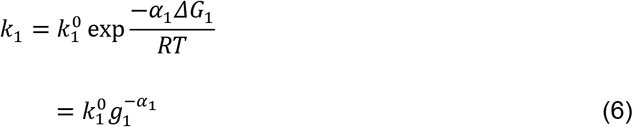

where 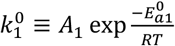 and 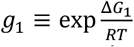 were used to aggregate factors independent and dependent on the driving force, respectively (see Supporting Information, Appendix 1 for details). *k*_1*r*_ and *k*_2_ can also be written similarly as:

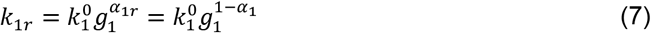

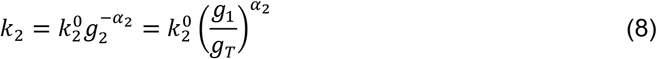

using notations similar to those defined for *k*_1_ (See Appendices 2 and 3 for details). Substituting these rate constants into Eq. (2) yields the following expression for *K*_*m*_:

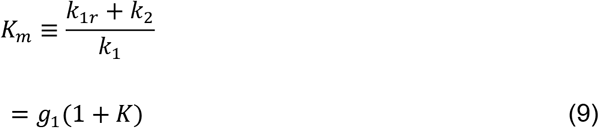

where *K* was defined as 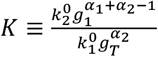 Finally, based on Eqs. (8) and (9), the enzymatic activity (*v*) can be expressed as:

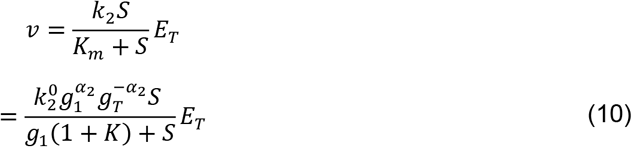

To illustrate how Eq. (10) captures the tradeoff relationship between *k*_2_ and *K*_*m*_, numerical simulations were performed (Fig. 2A). Hereafter, all simulations will be performed at α_1_ = α_1_*r* = α_2_ = 0.5, which is a common assumption used to make baseline models in heterogeneous catalysis.^[22,26-28]^ Physically, this means that when the driving force of an elementary reaction is increased by 1 kJ/mol, its activation barrier decreases by 0.5 kJ/mol. In reality, typical experimental values of α range between 0.3 and 0.7 for artificial catalysts,^[29-31]^ and the experimental value reported for cellulases is 0.74.^[25]^ Therefore, the influence of α deviating from 0.5 will be discussed in detail in Fig. 5D.

**Figure 2.**
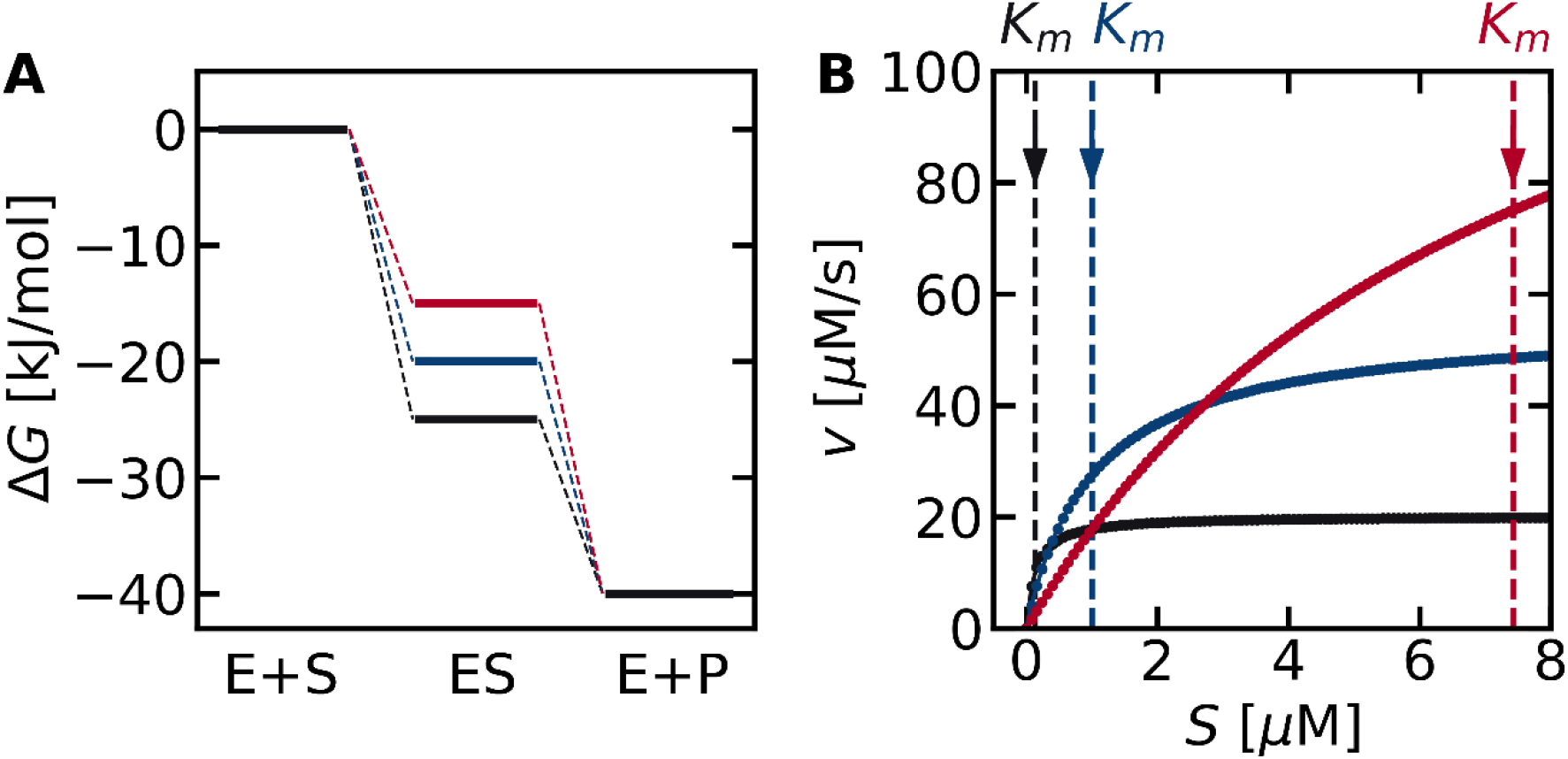
Thermodynamic landscapes (A) and their corresponding activity shown in the form of Michaelis-Menten plots (B). The *K*_*m*_ values are indicated as vertical dashed lines in (B). Increasing the driving force of the first step increases activity at low substrate concentrations but lowers the activity at high substrate concentrations. Therefore, the thermodynamic landscape of an optimum enzyme depends on the substrate concentration (*S*). The free energies of the enzyme-substrate complex (Δ*G*_1_) were −25, −20, and − 15 kJ/mol for the black, blue, and red lines, respectively, and that of the total reaction (Δ*G*_*T*_) was −40 kJ/mol. All numerical simulations in this study were performed at *E*_*T*_ = 1 µM, 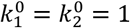 (1/*µM*/*s* and 1/s units, respectively).

Fig. 2A shows three possible thermodynamic landscapes for a reaction with a total driving force of Δ*GT* = −40 kJ/mol. This parameter was chosen as a representative value based on the fact that the Δ*GT* of typical biochemical reactions is between −80 ~ + 40 kJ/mol.^[32,33]^ Similar calculations with different values of Δ*GT* can be found in Figs. S1-S3. When the first reaction is thermodynamically favorable compared to the second (Δ*G*_1_ < Δ*G*_2_; Fig. 2A, black lines), the activity increases rapidly from low substrate concentrations (Fig. 2B, solid black line), consistent with the small *K*_*m*_ value. However, an enzyme with a small *K*_*m*_ suffers from a small *k*_2_ value, which is evident from the saturating behavior at *S* > 1 µM. Increasing the driving force of the second step (blue and red lines) leads to a larger *k*_2_ and thus higher activity at large *S* values (*S* > 1 µM) compared to the enzyme shown in black. At the same time, however, *K*_*m*_ increases, which decreases the enzymatic activity at low *S* (*S* < 1 µM). The difference in activity at low and high substrate concentrations occurs because the substrate participates in only the first elementary step. For example, even if *k*1 < *k*2 (Δ*G*1 > Δ*G*2), the rates of the two forward reactions (*k*1*E* ∙ *S* and *k*_2_(*ES*)) can be matched if the substrate concentration (*S*) is sufficiently large. However, at low substrate concentrations, a small *k*_1_ can no longer be compensated, resulting in the first step being rate-limiting. For this reason, a large *k*_1_ is necessary to increase the enzymatic activity at low substrate concentrations, whereas a large *k*_2_ is more desirable when the substrate concentration is sufficient. The balance in tradeoff changes when the rates of the two forward reactions are equal 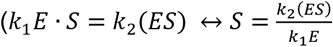 As the optimum values of *k*_1_ and *k*_2_ are dependent on the substrate concentration (*S*), the *K*_*m*_ value necessary to maximize the activity must also be dependent on (*S*).

### Analysis of the Activity – Driving Force Relationship

To directly illustrate the influence of driving force ( Δ*G*_1_ and Δ*GT* ) on enzymatic activity, we performed numerical simulations using Eq. (10) at various substrate concentrations (Fig. 3). At a substrate concentration of 0.1 *μ*M (Fig. 3A), the region of highest enzymatic activity (orange) was observed in the bottom left region. It is reasonable for activity to be higher in the lower half of the panel, due to the more negative Δ*GT*. A negative Δ*G*_1_ is also beneficial for activity at a low substrate concentration (*S* =0.1 *μ*), leading to enzymatic activity being higher in the left half of the panel. At higher substrate concentrations, the overall color within each panel changed from blue to red, because a higher substrate concentration always increases activity (Figs. 3B-3D). At the same time, the Δ*G*_1_ corresponding to maximum activity gradually shifted positively (black dashed lines). This finding is consistent with Fig. 2 which shows that a more positive Δ*G*_1_ is desirable when the substrate concentration is increased. In each panel, the location with the highest activity at a given Δ*GT* value is shown as a dashed black line. Notably, when the *K*_*m*_ value was calculated at the (Δ*G*_1_, Δ*GT*) values under the dashed line using Eq. (9), the obtained value was always equal to the substrate concentration *S* in each panel. In other words, the dashed line is not only the ridge of the volcano plot, but also the contour line showing *K*_*m*_ = *S* . This suggests that the condition for maximizing enzymatic activity can be represented by *K*_*m*_ = *S*.

**Figure 3.**
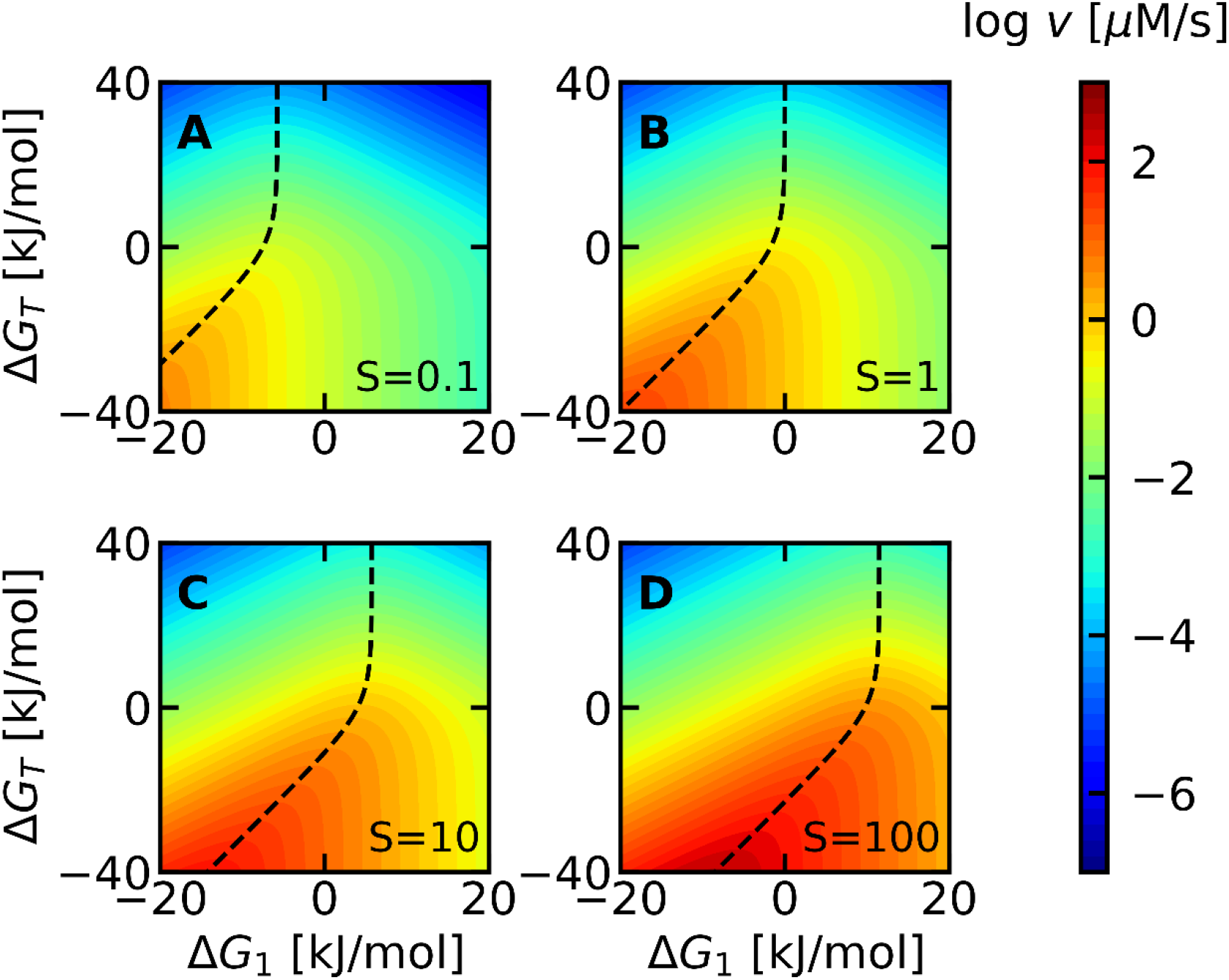
Enzymatic activity (*v*) plotted against Δ*G*_1_ and Δ*G*_*T*_ based on Eq. (10). The substrate concentration (S) in each panel was (A) 10^−1^, (B) 1, (C) 10, and (D) 10^2^ μM, as indicated in the bottom right of each panel. In all panels, the black dashed line corresponding to *K*_*m*_ = *S* overlaps with the region with the highest enzyme activity.

To examine why *K*_*m*_ = *S* leads to maximum activity, Eq. (10) was rearranged to give the following expression for the activity (*v*):

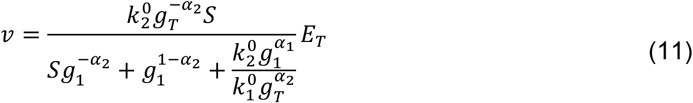

in which *g*_1_ is only in the denominator. The derivative of the denominator, denoted as *f* is:

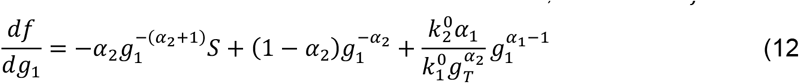

To maximize the activity (*v*), *f* must be minimized which is realized at:

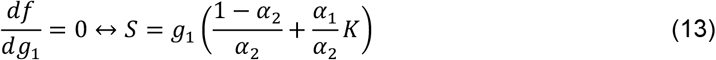

Considering that *K*_*m*_ is defined as *K*_*m*_ ≡ *g*_1_(1 + *K*) (Eq. (10), Eq. (13) yields a surprisingly simple formula for the condition of maximum activity when α1 = α1*r* = α2 = 0.5:

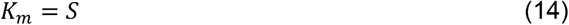

Eq. (14) provides the theoretical basis for why maximum activity was consistently observed along the contour line *K*_*m*_ = *S* in Fig. 3: The combination of (Δ*G*_1_, Δ*GT*) necessary to maximize activity guarantees *K*_*m*_ = *S*. This finding is further illustrated in Fig. 4, where the activity (*v*) is plotted as a function of *K*_*m*_ at different substrate concentrations. In all cases, maximum activity (*v*) is observed when the binding affinity (*K*_*m*_) is equal to the substrate concentration (*S*). Thus, the derivations and simulations so far provide mathematical evidence that having a *K*_*m*_ value equal to the substrate concentration *S* guarantees maximal enzymatic activity as long as the enzyme follows the Michaelis-Menten mechanism (Scheme 1), and the rate constants follow the BEP relationship with *α*_1_ = *α*_1*r*_ = *α*_2_ = 0.5.

**Figure 4.**
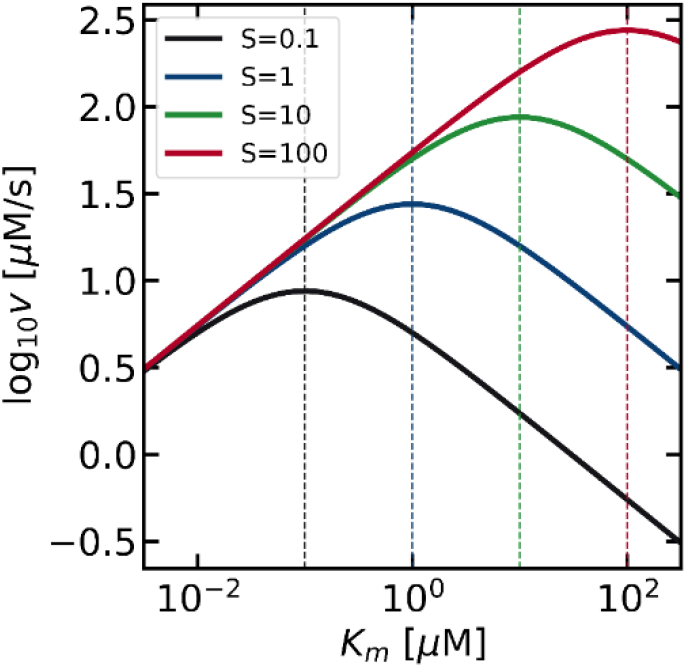
Volcano plots showing how the activity is expected to change with respect to the Michaelis-Menten constant ( *K*_*m*_ ). As the substrate concentration was increased from 10^−1^ μM (black) to 10^2^ μM (red), the volcano plot shifted to the upper right. The apex is located at *K*_*m*_ = *S*, as indicated by the vertical dashed lines of the corresponding color. Δ*G*_*T*_ = −40 kJ/mol and 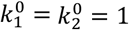 [1/μM/s and 1/s units, respectively] were used for the numerical simulations. Changing these values did not influence the conclusion that the activity is maximized when *K*_*m*_ = *S*, as shown in Fig. S4.

### Robustness of the Theoretical Model

To confirm the robustness of our finding, we have performed numerical simulations by loosening each of the theoretical requirements. Deviation from the Michaelis-Menten mechanism (Scheme 1) are shown in Fig. 5A-C, and deviation of α values from 0.5 are shown in Fig. 5D. The possibility of reverse reactions (P→S) or inhibition (E + I → EI or ES + I → ESI) are common deviations from Michaelis-Menten kinetics.^[34]^ The net rate in the presence of a reverse reaction when the substrate and product are in equal concentrations (*S* = *P* = 10 µM) is shown in Fig. 5A. In terms of maximizing the activity in the forward direction (S → P), the physically meaningful region is (Δ*GT* < 0), where the net reaction proceeds in the forward direction. Under this condition, the dashed line corresponding to *K*_*m*_ = *S* and the solid line corresponding to the true maximum activity (forward minus reverse reaction rates) overlap almost completely, indicating that *K*_*m*_ = *S* is a good guideline to enhance activity even in the presence of reverse reactions (P → S).

**Figure 5.**
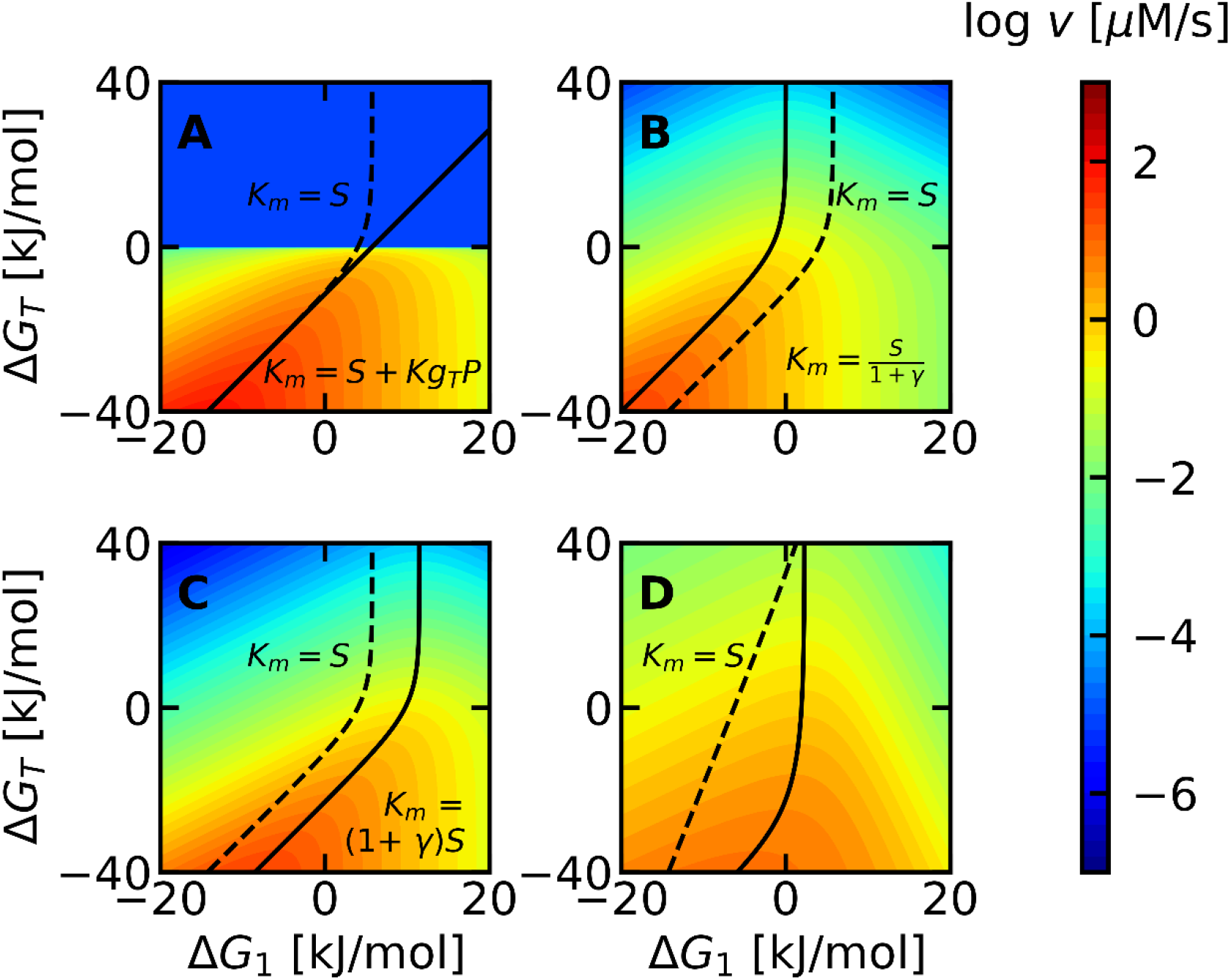
Influence of (A) Reverse reaction, (B) Competitive inhibition, (C) Uncompetitive inhibition, and (D) BEP coefficient, α on the optimal *K*_*m*_. The dashed line corresponds to *K*_*m*_ = *S*, with *S* = 10 µM. The true optimum *K*_*m*_ for each mechanism is shown as a solid line along with its analytical equation (refer to the SI for the derivations). In panel A, the product concentration (*P*) was set to 10 µM. The top half of (A) was colored at an arbitrarily low activity because the reverse reaction is more favorable in this region. The large discrepancy between the dashed and solid lines at Δ*G*_*T*_ > 0 is physically irrelevant, because the activity of the forward reaction cannot be discussed when the net reaction proceeds in the reverse direction. In panels B and C, the degree of inhibition (*γ* ≡ *I*/*K*_*i*_) was set to 10. In panel D, the BEP coefficients were set to *α*_1_ = *α*_2_ = 0.2. No analytical solution was obtained for D.

Similar calculations for competitive and uncompetitive inhibition, where the inhibitor binds to either the free enzyme or the enzyme-substrate complex, are shown in Fig. 5B,C. The degree of inhibition 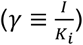, is determined by the inhibitor concentration (*I*) and the equilibrium constant of inhibition (*Ki*).^[34]^ Based on the experimental data of Park et al.,^[35]^ γ can range from 10^−4^ to 10^4^. As γ was less than 10 in approximately 80% of their data, γ = 10 was used here for the numerical simulations. Again, the optimal *K*_*m*_ (solid line) deviates only slightly from the dashed line (*K*_*m*_ = *S*), and both lines pass through the region of high activity (orange). The *K*_*m*_ values are approximately 1 order of magnitude apart between dashed and solid lines, yet there is only a 57 % difference in activity at a specific Δ*GT*. This is much smaller than the scale of the entire diagram (10 orders of magnitude), suggesting that adjusting *K*_*m*_ to the substrate concentration *S* is a robust strategy to enhance the activity, even in the presence of inhibition. A detailed discussion on the parameter dependence (γ, *S*), as well as for other mechanisms such as substrate inhibition or allostericity can be found in Section 3 of the supporting information. The derivations for the equations of the true optimal *K*^m^ can also be found in the same section.

The influence of the second assumption (α1 = α1*r* = α2 = 0.5) is shown in Fig. 5D. As physical constraints require α1*r* = 1 − α1 (Appendix 2), only α1 and α2 are independent. In an extreme case where α_1_ = α_2_ = 0.2, the activity is markedly diminished because rate constants hardly change even if their driving force is increased. However, the dashed line still passes through the region of high activity, and the activity is still less than an order of magnitude away from the true optimum (solid lines). Taken together, these simulations confirm that *K*_*m*_ = *S* is a robust theoretical guideline to enhance enzymatic activity.

### Validation based on Experimental Data

Finally, to evaluate whether *K*_*m*_ = *S* can rationalize enzymatic properties in nature, we have analyzed their relationship based on the experimental data from Park et al.^[35]^ The original data consisted of *K*_*m*_ values of wild-type enzymes obtained from BRENDA, and intracellular *S* values obtained from *Escherichia coli, Mus musculus*, and *Saccharomyces cerevisiae* cells, yielding a total of 1703 *K*_*m*_–*S* combinations. This dataset was then classified based on the number of entries for each substrate, based on the expectation that a substrate which participates in many reactions is more likely to deviate from Michaelis-Menten kinetics. ATP is the most frequent substrate with 313 entries and is shown in black. Both the raw *K*_*m*_ and *S* values (Fig. 6A) and their relative distribution (Fig. 6B) shows that *S* > *K*_*m*_ for ATP. The deviation from *K*_*m*_ = *S* may be because the Michaelis-Menten mechanism, which is the basis of our mathematical analysis, does not consider scenarios where multiple reactions compete for the same substrate. The next subset shown in blue covers 410 entries and consists of 5 substrates which each appear more than 50 times: NAD^+^, NADH, NADP^+^, NADPH, and acetyl-CoA. These cofactors are less universal than ATP, and *S* is only slightly larger than *K*_*m*_. The remaining 980 entries are shown in red. This subset contains 115 substrates such as carbon metabolites and amino acids and appear within the dataset 8 times on average. As the substrate becomes less universal, their *K*_*m*_ and *S* values become roughly consistent. In particular, the Gaussian distribution fitted to the red histogram (Fig. 6B) has a center at log_10_ *S*/*K*_*m*_ = 0.18 and a standard deviation of 1.3, which is reasonable considering that influences from inhibitors or the BEP coefficient can change the optimum *K*_*m*_ by roughly an order of magnitude (Fig. 5). Thus, the dataset from wild-type enzymes supports the theoretical prediction that a Michaelis-Menten constant equivalent to the substrate concentration is favorable for the activity, especially when the substrate participates in fewer reactions and Michaelis-Menten kinetics becomes more accurate.

**Figure 6.**
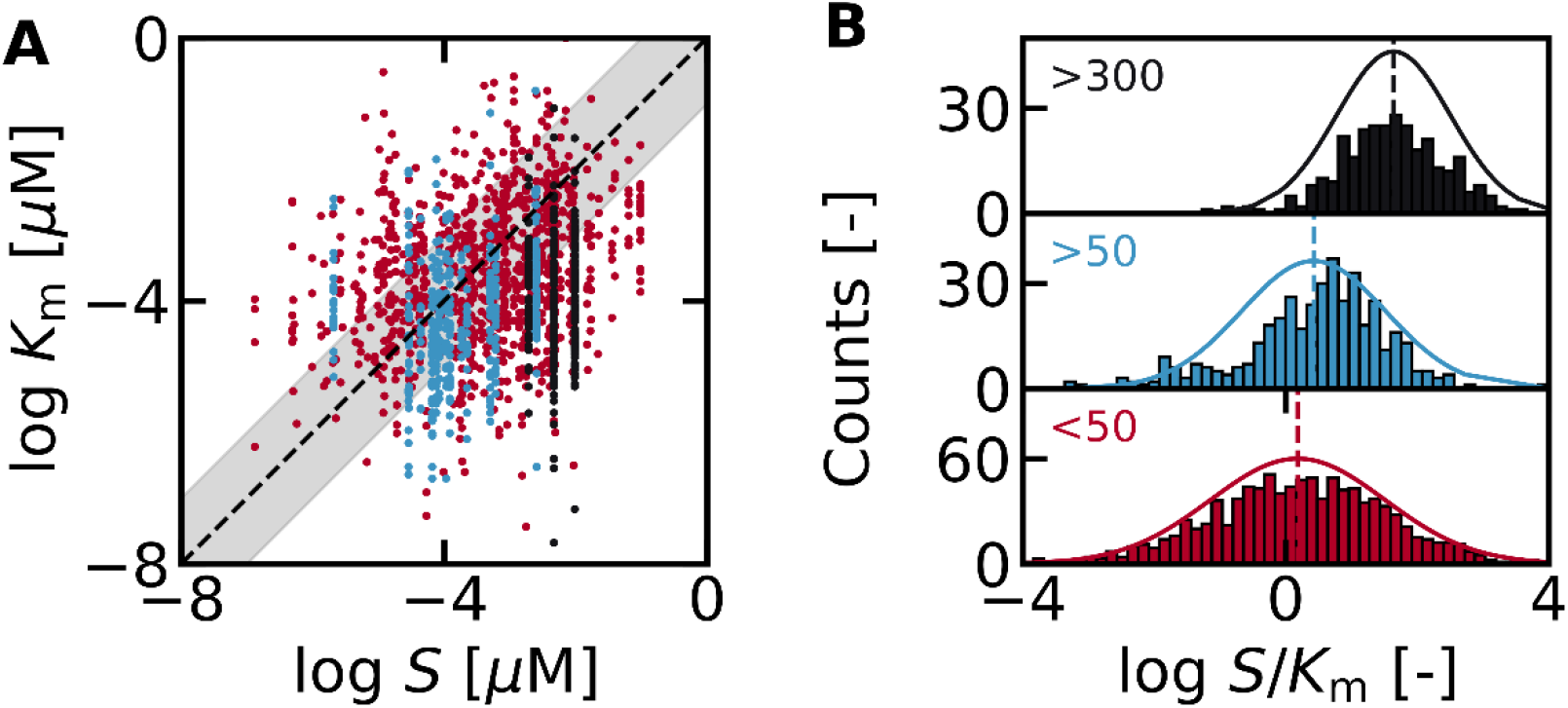
Relationship between *K*_*m*_ and *S* from the dataset reported by Park et al.^[35]^ The raw values of *K*_*m*_ and *S* are shown in (A), and their relative values are plotted in (B). Each entry of *K*_*m*_ and *S* was categorized based on the number of times the substrate appeared in the entire dataset. Black: > 300 (ATP), blue: > 50 (NAD^+^, NADH, NADP^+^, NADPH, and acetyl-CoA), red: < 50 (others). The number of entries was used as a proxy for the validity of the Michaelis-Menten mechanism of the specific substrate. The dashed line in (A) corresponds to *K*_*m*_ = *S*, and the shaded area shows a deviation of 1 order of magnitude.

## Discussion

So far, various criteria^[13,34,36]^ such as large *k*2 ( *kcat* ), small *K*_*m*_, or large *k*2/*K*_*m*_ have been proposed to characterize enzymes with high activity, making it difficult to rationally evaluate or engineer the activity of an enzyme. The lack of a universal consensus is largely due to the mutual dependence between *k*_2_ and *K*_*m*_. As our theoretical model addresses this challenge directly and maximizes the activity within the thermodynamic constraints imposed by *k*_2_ and *K*_*m*_, we believe that *K*_*m*_ = *S* is a criterion for high activity which is viable in a wider range of scenarios.

The idea that the Michaelis-Menten constant should be increased at higher substrate concentrations to maximize activity is consistent with the experimental work by Kari et al,^[37]^ who measured the activity of cellulases with different *K*_*m*_ . When the substrate concentration was increased 6 times, the *K*_*m*_ value of the most active enzyme increased approximately 2.4 times. Considering that *K*_*m*_ can change by roughly 6 orders of magnitude, the experimental trend supports our hypothesis *K*_*m*_ = *S*, especially when their experimental BEP coefficient of 0.74 is considered. The idea of the optimum binding affinity being dependent on the reaction condition and driving force is also consistent with recent theoretical models of heterogeneous catalysis.^[22,38-40]^

As a corollary, our model which quantifies the relationship between *K*_*m*_ and *k*_2_ immediately rationalizes the recently reported free energy relationship between them in cellulases.^[25]^ Namely, the relationship between *K*_*m*_ and *k*_2_ can be written as:

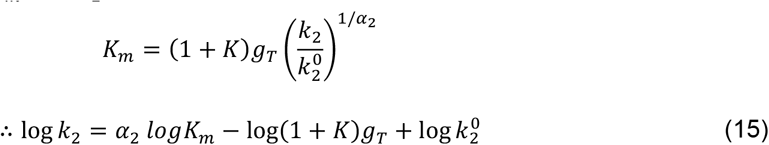

This equation shows that log *k*_2_ and log *K*_*m*_ are linearly correlated by a factor of α_2_, and provides a physical basis to the high linearity (R_2_ = 0.95) observed for cellulases.^[25]^ The consistency between our theoretical model and previously accumulated experimental insight suggests that it may be possible to quantitatively rationalize enzymatic properties based on fundamental principles of physical chemistry.

### Online Methods

The mathematical formulas were derived by hand, and the step-by-step derivations for the standard Michaelis-Menten mechanism are explained in the main text. The derivations in the presence of inhibition and allostericity are provided in the supporting information. Numerical simulations and bioinformatic analysis were performed using Python 3.9.12. The code used for the analysis can be found in the extended data or accessed directly at github: https://github.com/HideshiOoka/SI_for_Publications.

## Supporting information

Main SI

## Acknowledgments

H.O. gratefully acknowledges the support from the JST FOREST program (Grant Number JPMJFR213E, Japan). Y. C. is grateful for the support from the JST ACT-X program (Grant Number JPMJAX20BB, Japan).

## Notes

### Competing Interest Statement

The authors have declared no competing interest.

