## Supplementary material for "Universal Design Principle to Enhance Enzymatic Activity using the Substrate Affinity": Main SI

### Supporting Information for: Universal Design Principle to Enhance Enzymatic Activity using the Substrate Affinity

†*Biofunctional Catalyst Research Team*

*RIKEN Center for Sustainable Resource Science (CSRS)*

*2-1 Hirosawa, Wako, Saitama 351-0198, Japan*

‡ *Earth-Life Science Institute (ELSI), Tokyo Institute of Technology*

*2-12-IE-1, Ookayama, Meguro-ku, Tokyo 152-8550, Japan*

**2 Influence of the Driving Force ($\Delta G_T$)** **6**

**3 Influence of the Rate Constants $k_1^0$ and $k_2^0$** **8**

**4 Deviations from Michaelis-Menten Kinetics** **10**

### 1 Mathematical Details

#### Appendix 1.

The mathematical details to obtain Eq. (6) in the main text is shown below. Based on Eq. (4), the activation barrier ($E_{a1}$) can be written as:

$$E_{a1} = E_{a1}^0 + \alpha_1 \Delta G_1 \tag{S1}$$

Using this to substitute $E_{a1}$ in Eq. (5) yields:

$$k_1 = A_1 \exp \frac{-E_{a1}}{RT} \tag{S2}$$
$$= A_1 \exp \frac{-E_{a1}^0 - \alpha_1 \Delta G_1}{RT} \tag{S3}$$
$$= A_1 \exp \frac{-E_{a1}^0}{RT} \exp \frac{-\alpha_1 \Delta G_1}{RT} \tag{S4}$$
$$\tag{S5}$$

By grouping the factors independent of $\Delta G_1$ as follows:

$$k_1^0 \equiv A_1 \exp \frac{-E_{a1}^0}{RT} \tag{S6}$$

Eq. (6) in the main text can be obtained:

$$k_1 = k_1^0 \exp \frac{-\alpha_1 \Delta G_1}{RT} \tag{S7}$$

#### Appendix 2.

The mathematical details to obtain Eq. (7) are shown below. The activation barrier for $k_{1r}$ can be expressed in a similar way to Eq. (4) in the main text as:

$$E_{a1r} = E^0_{a1r} + \alpha_{1r}\Delta G_{1r} \tag{S8}$$

Considering that $\alpha_1 + \alpha_{1r} = 1$ and $\Delta G_1 = -\Delta G_{1r}$, $E_{a1r}$ can be expressed as:

$$E_{a1r} = E^0_{a1r} + (\alpha_1 - 1)\Delta G_1 \tag{S9}$$

Therefore, based on the Arrhenius equation, $k_{1r}$ can be expressed as:

$$k_{1r} = A_{1r} \exp \frac{-E_{a1r}}{RT} \tag{S10}$$

$$= A_{1r} \exp \frac{-E^0_{a1r} - (\alpha_1 - 1)\Delta G_1}{RT} \tag{S11}$$

$$= A_{1r} \exp \frac{-E^0_{a1r}}{RT} \exp \frac{(1 - \alpha_1)\Delta G_1}{RT} \tag{S12}$$

$$= k^0_1 g_1^{1-\alpha_1} \tag{S13}$$

This is Eq. (7) in the main text.

#### Appendix 3.

The mathematical details to obtain Eq. (8) are shown below. The activation barrier for $k_2$ can be expressed in a similar way to Eq. (4) in the main text as:

$$E_{a2} = E_{a2}^0 + \alpha_2 \Delta G_2 \tag{S14}$$

Therefore, based on the Arrhenius equation, $k_2$ can be expressed as:

$$\begin{aligned} k_2 &= A_2 \exp \frac{-E_{a2}}{RT} && \text{(S15)} \\ &= A_2 \exp \frac{-E_{a2}^0 - \alpha_2 \Delta G_2}{RT} && \text{(S16)} \\ &= A_2 \exp \frac{-E_{a2}^0}{RT} \exp \frac{-\alpha_2 \Delta G_1}{RT} && \text{(S17)} \\ &= k_2^0 g_2^{-\alpha_2} && \text{(S18)} \end{aligned}$$

Taking into account $\Delta G_2 = \Delta G_T - \Delta G_1$, $g_2 = g_T/g_1$. Therefore,

$$k_2 = k_2^0 \left( \frac{g_1}{g_T} \right)^{\alpha_2} \tag{S19}$$

where $k_2^0$ was defined in a way similar to $k_1^0$ in the main text, namely $k_2^0 \equiv A_2 \exp \frac{-E_{a2}^0}{RT}$. This corresponds to Eq. (8) in the main text.

#### 2 Influence of the Driving Force ($\Delta G_T$)

Numerical simulations of Michaelis-Menten kinetics at varying driving forces ($\Delta G_T$) are provided in this section. The python code provided as part of the supplementary information can be used to generate similar plots with different parameter values.

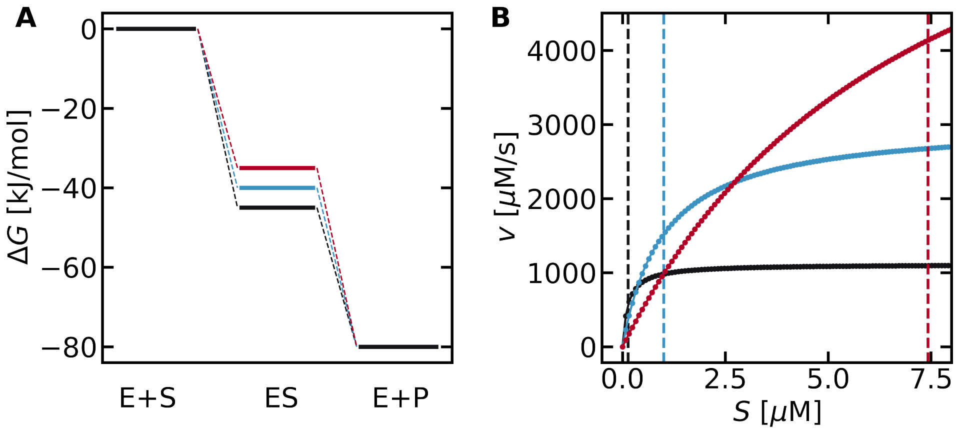

Figure S1: Influence of the driving force at $\Delta G_T = -80$ kJ/mol and $\Delta G_1 = -35, -40, -45$ kJ/mol. All other parameters are the same as in Fig. 2 of the main text.

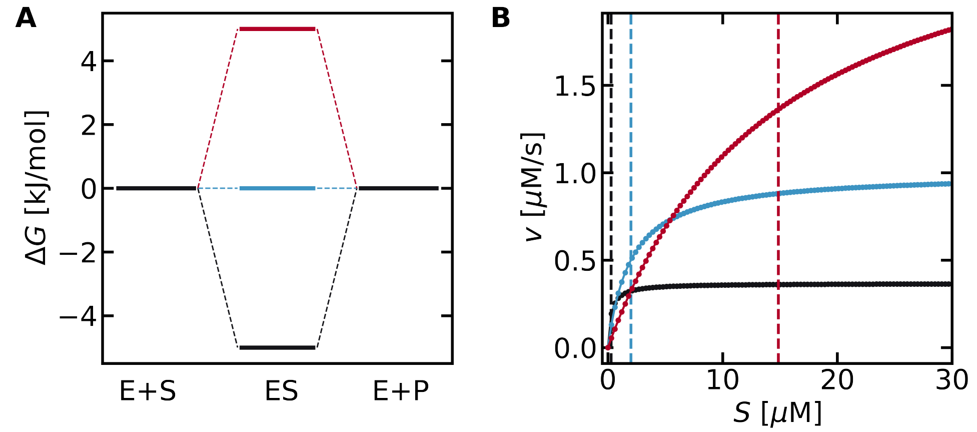

Figure S2: Influence of the driving force at $\Delta G_T = 0$ kJ/mol and $\Delta G_1 = -5, 0, +5$ kJ/mol. All other parameters are the same as in Fig. 2 of the main text.

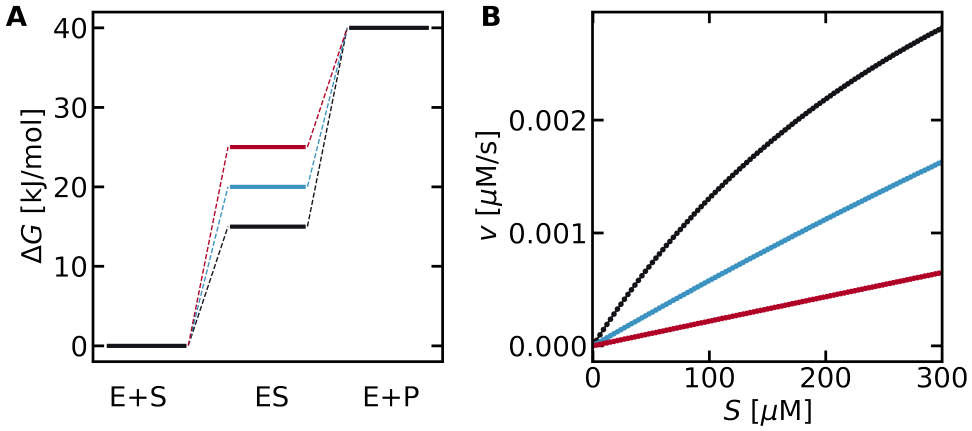

Figure S3: Influence of the driving force at $\Delta G_T = 40$ kJ/mol and $\Delta G_1 = 15, 20, 25$ kJ/mol. All other parameters are the same as in Fig. 2 of the main text.

### 3 Influence of the Rate Constants $k_1^0$ and $k_2^0$

The values $k_1^0 = k_2^0 = 1$ were used in the main text, due to a necessity to set parameter values for the numerical simulations. However, the values of $k_1^0$ and $k_2^0$ do not influence the mathematical conclusion that the activity $v$ can be maximized at $K_m = S$, as can be seen in the simulations below.

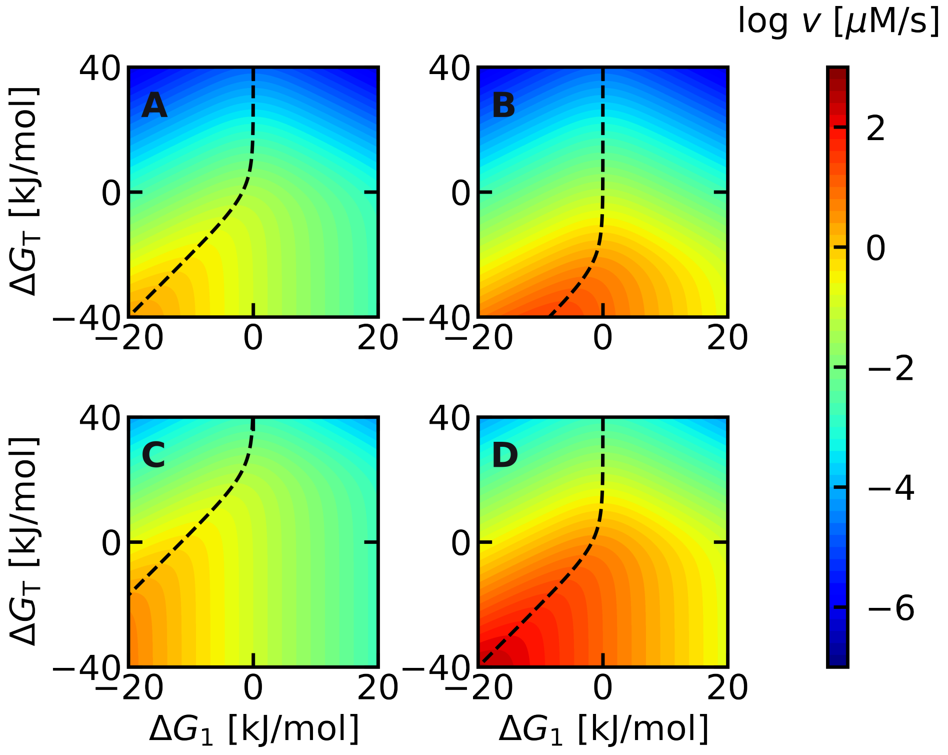

Figure S4: Influence of $k_1^0$ and $k_2^0$ on the location of the optimum $K_m$. $(k_1^0, k_2^0)$ = (0.1,0.1), (10,0.1),(0.1,10),(10,10) in panels A-D. The substrate concentration was set to 1. Regardless of the values of $k_1^0$ and $k_2^0$, maximum activity is observed at $K_m = S$ (dashed line).

As explained in the main text, the essential requirement to achieve maximum activity is that the derivative of Eq. (13) becomes zero. This is achieved when

$$g_1(1 + K) = S \tag{S20}$$

As $K_m \equiv g_1(1+K)$ (Eq. 9), the condition $K_m = S$ guarantees that the derivative of Eq. (13)

becomes zero regardless of other parameters such as $g_1, g_T, \Delta G_1, \Delta G_T, k_1^0, k_2^0, K$. Within the scope of the model presented in the main text, the only way that the optimum $K_m$ deviates from $S$ is to break the assumptions leading up to Eq. (14), which is the Michaelis-Menten mechanism, and $\alpha_1 = \alpha_{1r} = \alpha_2 = 0.5$, both of which were tested directly in Fig. 5.

### 4 Deviations from Michaelis-Menten Kinetics

The rate laws of various chemical mechanisms will be derived in this section. We will begin our evaluation of starting from the original Michaelis-Menten equation and then evaluate how the various mechanisms will modify the rate laws.

#### 4.1 Michaelis-Menten Kinetics

The standard Michaelis-Menten mechanism is given as follows:

$$\mathrm{E} + \mathrm{S} \underset{k_{1r}}{\overset{k_1}{\rightleftharpoons}} \mathrm{ES} \xrightarrow{k_2} \mathrm{E} + \mathrm{P} \tag{S21}$$

The time dependence of the concentration of ES can be expressed as:

$$\frac{dES}{dt} = k_1 E \cdot S - (k_{1r} + k_2)ES \tag{S22}$$

Hereafter, chemical species (E,S,ES,P) will be written in upright font, and their concentrations ($E, S, ES, P$) will be expressed in italics. Using these notations, the steady-state concentration of ES can be expressed as:

$$ES = \frac{k_1 S}{k_{1r} + k_2} E \tag{S23}$$

$$= \frac{S}{K_m} E \tag{S24}$$

where the definition of $K_m \equiv \frac{k_{1r}+k_2}{k_1}$ was used in the last step. As the total enzyme concentration ($E_T$) must be constant, we obtain:

$$E_T = E + ES \tag{S25}$$
$$= (1 + \frac{S}{K_m})E \tag{S26}$$
$$= \frac{S + K_m}{K_m}E \tag{S27}$$

Therefore,

$$E = \frac{K_m}{S + K_m}E_T \tag{S28}$$
$$ES = \frac{S}{S + K_m}E_T \tag{S29}$$

This yields the final rate as:

$$v = k_2 ES \tag{S30}$$
$$= \frac{k_2 S}{K_m + S}E_T \tag{S31}$$

#### 4.2 Reverse Reactions

Suppose the reverse reaction cannot be ignored ($k_{2r}P > 0$). In that case, an additional term ($k_{1r}k_{2r}P$) must be added to the numerator to the rate law in the main text to yield:

$$v = \frac{k_1 k_2 S - k_{1r} k_{2r} P}{k_1 S + k_{1r} + k_2 + k_{2r} P} E_T \tag{S32}$$
$$= \frac{k_1^0 k_2^0 / \sqrt{g_T} S - k_1^0 k_2^0 \sqrt{g_T} P}{k_1^0 / \sqrt{g_T} S + k_1^0 \sqrt{g_1} + k_2^0 \sqrt{\frac{g_1}{g_T}} + k_2^0 \sqrt{\frac{g_T}{g_1}} P} E_T \tag{S33}$$
$$= \frac{k_2^0}{\sqrt{g_T}} \frac{S - g_T P}{(S + K g_T P)/\sqrt{g_1} + (1 + K)\sqrt{(g_1)}} \tag{S34}$$

Setting the denominator as $f$, we have

$$\frac{df}{dg_1} = -0.5g_1^{-1.5}(S + Kg_TP) + 0.5g_1^{-0.5}(1 + K) = 0 \tag{S35}$$

$$\Leftrightarrow g_1(1 + K) = S + Kg_TP \tag{S36}$$

indicating that $K_m = S + Kg_TP$ is the condition for maximum activity. When the forward reaction is favored ($\Delta G_T < 0 \leftrightarrow g_T \ll 1$), the optimum $K_m$ converges the the substrate concentration $S$.

#### 4.3 Competitive Inhibition

Competitive inhibition occurs when an inhibitor competes for the active site of the enzyme via the following reaction:

$$\mathrm{E} + \mathrm{I} \underset{k_{ir}}{\overset{k_i}{\rightleftharpoons}} \mathrm{EI} \tag{S37}$$

Let $K_i \equiv k_{ir}/k_i$ denote the extend of inhibition. Then, the steady-state concentration of EI can be expressed as:

$$EI = \frac{I}{K_i}E \tag{S38}$$

$$= \gamma E \tag{S39}$$

where we have set $\gamma \equiv \frac{I}{K_i}$. In this case, the total enzyme concentration can be expressed as:

$$ET = E + ES + EI \tag{S40}$$

$$= (1 + \gamma)E + ES \tag{S41}$$

$$= (1 + \gamma + \frac{S}{K_m})E \tag{S42}$$

Note that the steady-state concentration of ES was expressed using Eq. S23. Based on the above, E and ES can be expressed as follows:

$$E = \frac{(1+\gamma)K_m}{(1+\gamma)K_m + S} E_T \tag{S43}$$

$$ES = \frac{S}{(1+\gamma)K_m + S} E_T \tag{S44}$$

Therefore, the enzymatic activity can be expressed as:

$$v = k_2 ES \tag{S45}$$

$$= \frac{k_2 S}{(1+\gamma)K_m + S} E_T \tag{S46}$$

This rate expression is similar to the case without inhibition (Eq. S31). The only difference is that there is an extra term $(1+\gamma)$ before $K_m$, which will shift the optimum value of $K_m$ to $S' \equiv S/(1+\gamma)$

In order to compare the influence of competetive inhibition, let us calculate the relative rates at $K_m = S$ and $K_m = S'$. In the case of $K_m = S$,

$$v_{K_m=S} = \frac{k_2 S'}{S + S'} E_T \tag{S47}$$

$$= \frac{k_2(1+\gamma)}{2+\gamma} E_T \tag{S48}$$

$$= \frac{k_2(1+\gamma)}{2+\gamma} E_T \tag{S49}$$

On the other hand, in the case of $K_m = S'$

$$v_{K_m=S'} = \frac{k_2 S'}{S' + S'} E_T \tag{S50}$$

$$= \frac{1}{2} k_2 E_T \tag{S51}$$

It is tempting to divide Eq. S51 with Eq. S49 directly to obtain a relative activity of

$$\frac{v_{K_m=S}}{v_{K_M=S'}} = \frac{2(1+\gamma)}{2+\gamma} \tag{S52}$$

However, this approach is flawed. For example, Eq. S52 suggests that at strong inhibition ($\gamma \to \infty$), $\frac{v_{K_m=S}}{v_{K_M=S'}} > 1$. This cannot be correct, as the activity should be maximized at $K_m = S'$. The main problem is that $k_2$ is dependent on $K_m$, and therefore, the $k_2$ in Eq. S51 and Eq. S49 should not have been canceled out.

In order to take the relationship between $K_m$ and $k_2$ into consideration, we have used the notations in the main text ($K_m = g_1(1+K), k_2 = k_2^0\sqrt{\frac{g_1}{g_T}}$). When $K_m = S$, $g_1 = \frac{S}{1+K}$. Therefore, Eq. S49 becomes

$$v_{K_m=S} = \frac{1+\gamma}{2+\gamma} k_2^0 \sqrt{\frac{S}{(1+K)g_T}} E_T \tag{S53}$$

Similarly, when $K_m = S'$, $g_1 = \frac{S'}{1+K}$. Therefore, Eq. S51 becomes

$$v_{K_m=S'} = \frac{1}{2} k_2^0 \sqrt{\frac{S'}{(1+K)g_T}} E_T \tag{S54}$$

Finally, dividing Eq. S54 with Eq. S53 gives the following expression of relative activity:

$$\frac{v_{K_m=S}}{v_{K_m=S'}} = \frac{2(1+\gamma)}{2+\gamma}\sqrt{\frac{S}{S'}} \tag{S55}$$

$$= \frac{2(1+\gamma)}{2+\gamma}\sqrt{\frac{1}{1+\gamma}} \tag{S56}$$

$$= \frac{2\sqrt{1+\gamma}}{2+\gamma} \tag{S57}$$

Note that based on the relationship between the arithmetic and geometric mean,

$$\frac{1 + (1 + \gamma)}{2} \geq \sqrt{1 \cdot (1 + \gamma)} \tag{S58}$$

$$\tag{S59}$$

Therefore,

$$\frac{v_{K_m=S}}{v_{K_m=S'}} = \frac{2\sqrt{1+\gamma}}{2+\gamma} \leq 1 \tag{S60}$$

showing that $K_m = S'$ is indeed the optimum value of $K_m$ under competitive inhibition. The relative activity when $\gamma$ is varied is shown below.

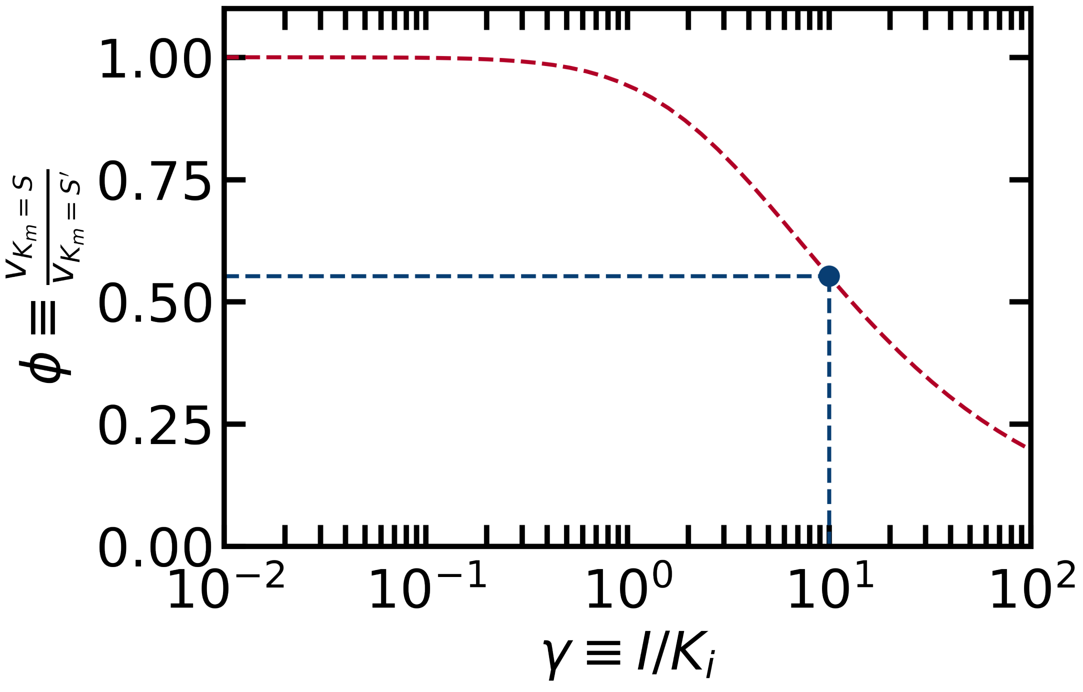

Figure S5: Influence of competitive inhibition for $\gamma$ between $10^{-3}$ and $10^{3}$. Fig. 5B in the main text corresponds to the condition where $\gamma = 10$ (blue circle).

#### 4.4 Uncompetitive Inhibition

In the case of uncompetitive inhibition, the following reaction decreases the concentration of the active enzyme:

$$\text{ES} + \text{I} \underset{k_{ir}}{\overset{k_i}{\rightleftharpoons}} \text{ESI} \tag{S61}$$

As in the case of competitive inhibition, let $K_i \equiv k_{ir}/k_i$. Then, the steady-state concentration of ESI can be expressed as:

$$\text{ESI} = \frac{I}{K_i} ES \tag{S62}$$

$$= \gamma ES \tag{S63}$$

where we have set $\gamma \equiv \frac{I}{K_i}$. In this case, the total enzyme concentration can be expressed as:

$$ET = E + ES + ESI \tag{S64}$$

$$= E + (1+\gamma)ES \tag{S65}$$

$$= (1 + \frac{S}{K_m}(1+\gamma))E \tag{S66}$$

$$E_T = E + ES + ES_2 \tag{S67}$$

$$= E + (1+\gamma)ES \tag{S68}$$

$$= (1 + \frac{S}{K_m}(1+\gamma))E \tag{S69}$$

Therefore,

$$E = \frac{K_m}{(1+\gamma)S + K_m} E_T \tag{S70}$$

$$ES = \frac{(1+\gamma)S}{(1+\gamma)S + K_m} E_T \tag{S71}$$

This yields the final rate as:

$$v = k_2 ES \tag{S72}$$

$$= \frac{k_2(1+\gamma)S}{(1+\gamma)S + K_m} E_T \tag{S73}$$

$$= \frac{k_2 S'}{K_m + S'} E_T \tag{S74}$$

where the final expression was obtained by setting $S' \equiv (1+\gamma)S$.

This expression is mathematically equivalent to the rate expression without inhibition (Eq. S31) except that $S'$ is used in place of $S$. As the derivations in the main text show that the maximum rate of Eq. S31 is given by $K_m = S$, the maximum rate for Eq. S89 is given by $K_m = S'$. Therefore, the maximum rate in the case of uncompetive inhibition is given by:

$$K_m = S' \tag{S75}$$

$$= (1+\gamma)S \tag{S76}$$

The dependence on $\gamma$ has the exact same expression in the case of competitive inhibition (Eq. S60).

#### 4.5 Substrate Inhibition

In the case of substrate inhibition, the reaction is inhibited when the substrate binds excessively to the enzyme. For example, a second substrate molecule may bind to the enzyme-substrate complex ($ES_2$), according to the following reaction:

$$\mathrm{ES} + \mathrm{S} \underset{k_{ir}}{\overset{k_i}{\rightleftharpoons}} \mathrm{ES}_2 \tag{S77}$$

When $ES_2$ is in steady-state, the following equation holds:

$$k_i ES \cdot S = k_{ir} ES_2 \tag{S78}$$

$$\tag{S79}$$

Defining $K_i \equiv k_{ir}/k_i$ gives the following expression:

$$ES_2 = \frac{S}{K_i} ES \tag{S80}$$

$$= \gamma ES \tag{S81}$$

where $\gamma \equiv \frac{S}{K_i}$. As the total amount of the enzyme ($E_T$) must remain constant, the contribution from $ES_2$ effectively serves to decrease the concentration of ES which can be used to form the product. Namely,

$$E_T = E + ES + ES_2 \tag{S82}$$

$$= E + (1+\gamma)ES \tag{S83}$$

$$= (1 + \frac{S}{K_m}(1+\gamma))E \tag{S84}$$

Therefore,

$$E = \frac{K_m}{(1+\gamma)S + K_m} E_T \tag{S85}$$

$$ES = \frac{(1+\gamma)S}{(1+\gamma)S + K_m} E_T \tag{S86}$$

This yields the final rate as:

$$v = k_2 ES \tag{S87}$$

$$= \frac{k_2(1+\gamma)S}{(1+\gamma)S + K_m} E_T \tag{S88}$$

$$= \frac{k_2 S'}{K_m + S'} E_T \tag{S89}$$

where the final expression was obtained by setting $S' \equiv (1+\gamma)S$.

This expression is equivalent to competitive inhibition, because in both cases, an inhibitor decreases the concentration of ES by converting it into an inactive state. Substrate inhibition is a special case of competitive inhibition where the inhibitor is the substrate molecule itself. However, within the simplified framework of Michaelis-Menten kinetics, the identity of the inhibitor does not alter the final rate equation.

#### 4.6 Allostericity

As a final example of deviation from Michaelis-Menten kinetics, we have considered allostericity according to the following mechanism:

$$\mathrm{E + nS} \underset{k_{1r}}{\overset{k_1}{\rightleftharpoons}} \mathrm{ES_n} \xrightarrow{k_2} \mathrm{E + nP} \tag{S90}$$

This is a mechanism corresponding to positive homotropic allostericity such as in the case of hemoglobin, where the substrate is also an effector which enhances the activity.

Based on the steady-state approximation of $ES_n$, the following expression can be obtained:

$$k_1 E \cdot S^n = (k_{1r} + k_2) ES_n \tag{S91}$$

$$ES_n = \frac{k_1}{k_{1r} + k_2} ES^n \tag{S92}$$

$$= \frac{S^n}{K_m} E \tag{S93}$$

Eq. S93 is the same as Eq. S23 except that $S$ has been replaced with $S^n$. Therefore, the final rate equation can be obtained by replacing the $S$ in Eq. S31 with $S^n$:

$$v = \frac{k_2 S^n}{K_m + S^n} E_T \tag{S94}$$

This equation corresponds to the famous Hill-Langmuir equation of allosteric enzymes, and the optimum $K_m$ is given at $K_m = S^n$, not $K_m = S$. The difference in activity can be calculated from

$$v_{K_m=S} = \frac{k_2 S^n}{S + S^n} E_T \tag{S95}$$

$$= \frac{S^n}{S + S^n} \sqrt{\frac{S}{(1+K) g_T}} k_2^0 E_T \tag{S96}$$

and

$$v_{K_m=S^n} = \frac{k_2 S^n}{S^n + S^n} E_T \tag{S97}$$

$$= \frac{1}{2} k_2 E_T \tag{S98}$$

$$= \frac{1}{2} \sqrt{\frac{S^n}{(1+K) g_T}} k_2^0 E_T \tag{S99}$$

as

$$v_{K_m=S}/v_{K_m=S^n} = \frac{2S^{\frac{1+n}{2}}}{S+S^n} \tag{S100}$$

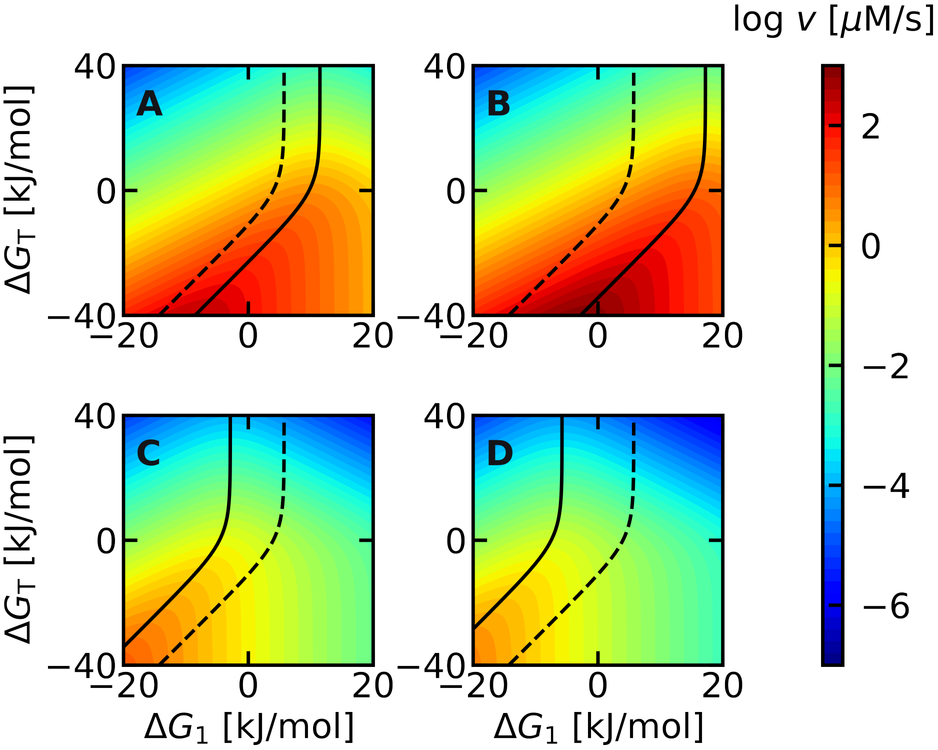

Figure S6: Influence of allostericity with a Hill constant ($n$) of (A) 2, (B) 3, (C) -0.5, (D) -1. Other parameters are the same as Fig. 5 in the main text. Even under varying degrees of allostericity, the dashed line ($K_m = S$) is located near the solid line showing the true optimum $K_m$ ($K_m = S^n$) and passes through the high activity region (orange).
